# Discovery of Selective Small-Molecule Ligands of SV2C by AI-Enhanced Virtual Screening and Experimental Validation

**DOI:** 10.64898/2026.08.11.744237

**Authors:** Alexander C. Brueckner, Matthew F. Martin, Sheenam Khuttan, Benjamin Shields, Anshumali Mittal, Jennifer A. Schreiber, Romelia Salomon-Ferrer, Andrea Bortolato, Ali Salahpour, Meghan L. Bucher, Jonathan A. Coleman, Gary W. Miller

**Affiliations:** SandboxAQ, Palo Alto, California, United States; Department of Structural Biology, University of Pittsburgh, Pittsburgh, Pennsylvania, United States; University of Toronto, Toronto, Ontario, Canada; Columbia University Irving Medical Center, New York, New York, United States

**Author notes:** Corresponding authors: Alexander C. Brueckner, Gary W. Miller. These authors contributed equally (Alexander C. Brueckner, Matthew F. Martin).

## Abstract

Synaptic vesicle glycoprotein 2C (SV2C) is enriched in dopaminergic neurons and implicated in Parkinson’s disease, but no selective small-molecule probes exist for it. Lacking a full-length SV2C structure, we built a homology model from SV2A cryo-EM templates and used molecular dynamics to characterize its conformational landscape. An AI-enhanced virtual screening pipeline, validated on a curated SV2A benchmark, was applied to 5.96 million commercial compounds, prioritizing 94 candidates for experimental testing. Of 71 compounds tested in an orthogonal biophysical assay cascade, 22 were active (31% hit rate), and five advanced to isoform-selectivity profiling. Compounds 36 and 56 emerged as leads, with SV2C *K*_*i*_ values of 24.6 *µ*M and 3.25 *µ*M and greater than 10-fold selectivity over SV2A. An unpublished SV2A cryo-EM structure independently confirmed the predicted binding mode. This AI-driven pipeline delivered selective SV2C ligands from a general chemical library, providing tools to probe SV2C biology in Parkinson’s disease.

## 1 Introduction

The synaptic vesicle glycoprotein 2 (SV2) family comprises three 12-transmembrane proteins, SV2A, SV2B, and SV2C, that regulate vesicular neurotransmitter storage and release and play key roles in synaptic physiology. SV2A is widely expressed in central nervous system and is the established target of FDA-approved antiepileptic drug levetiracetam (LEV) and its analogs[1], and also serves as a neuronal receptor for botulinum neurotoxin A[2]. In contrast, SV2C exhibits a more restricted expression pattern, being enriched in dopaminergic neurons of the basal ganglia and related structures, where it contributes to dopamine homeostasis and vesicular storage.[3, 4]

Genetic, biochemical, and pathological evidence implicates SV2C in Parkinson’s disease (PD) and related disorders. SV2C levels are reduced following dopamine neuron loss in mouse models of PD, and SV2C knockout mice show reduced striatal dopamine content and release, along with mild motor deficits.[3] SV2C co-immunoprecipitates with *α*-synuclein, a central component of Lewy body pathology, and its expression is dramatically disrupted in PD brains but not in Alzheimer’s disease, progressive supranuclear palsy, or multiple system atrophy.[3] Genetic studies have identified SV2C polymorphisms associated with PD risk and altered responses to dopaminergic therapies, further underscoring its relevance in disease.[5] Together, these findings suggest that pharmacological modulation of SV2C may offer a novel therapeutic route to modify dopaminergic function or disease progression.

Despite its links to disease, the pharmacology of SV2C remains underdeveloped. Most small-molecule tools currently available are either pan-SV2 ligands, such as padsevonil[6], or SV2A-selective ligands such as LEV, brivaracetam (BRV)[7], and plosaracetam (PRM; also known as SDI-118 and ABBV-552)[8], which were discovered and optimized for antiseizure activity. Although a selective SV2C radioligand (UCB-F) has recently been characterized[9, 10], selective small-molecule modulators of SV2C, particularly positive modulators that enhance vesicular dopamine storage and controlled release, have not been reported. Several factors contribute to this gap: the lack of a full-length experimental SV2C structure; the conformational flexibility typical of major facilitator superfamily (MFS) transporters; and the high sequence and structural homology among SV2 isoforms (61–64% sequence identity and ∼80% structural similarity), which complicates isoform-selective design.

To address these challenges, we initiated a collaboration between SandboxAQ, SPARK NS, the University of Pittsburgh, Columbia University, and the University of Toronto to identify novel SV2C-selective small molecules using a combined computational-experimental strategy. The project was organized around four main aims: (1) to develop and validate an SV2C homology model and dynamic conformational ensemble informed by SV2A cryo-EM structures and molecular dynamics (MD) and Gaussian accelerated MD (GaMD)[11] simulations; (2) to retrospectively validate an AI-enhanced virtual screening (VS) pipeline on a curated SV2A ligand benchmark; (3) to apply this pipeline prospectively to a large CNS-biased subset of the Mcule in-stock library; and (4) to experimentally test prioritized hits using a biophysical screening cascade, followed by binding affinity and isoform selectivity characterization.

This VS campaign yielded an unusually high primary hit rate (31%) and identified multiple chemotypes with low-micromolar SV2C affinity and favorable selectivity over SV2A and SV2B, including compounds 36 and 56.

In this paper, we describe the design and performance of the AI-enhanced VS funnel, the biophysical screening cascade, and the initial structure-activity and isoform-selectivity landscape of the resulting hits. Specifically, we address three key questions: (i) can an SV2A-derived homology model and AI-driven VS pipeline enrich for bona fide SV2C binders in the absence of a full-length structure; (ii) what is the chemical and functional diversity of the identified hits, including their effects on SV2C stability and competition with a pan-SV2 ligand; and (iii) to what extent can SV2C selectivity versus SV2A/B be achieved at the hit stage, and what structural hypotheses emerge to guide further optimization?

## 2 Results and Discussion

### 2.1 SV2C Homology Modeling and Conformational Analysis

At the outset of this work, no full-length experimental structure of SV2C was available. We therefore constructed homology models of SV2C based on recently reported SV2A cryo-EM structures in complex with the padsevonil analogue UCB-2500 (PDB 8UO9)[6] and brivaracetam (PDB 8K77)[12]. Sequence alignment confirmed high conservation across the transmembrane core (61–64% sequence identity; ∼80% structural similarity), while highlighting localized differences near the putative racetam binding pocket and luminal domains, including SV2C Leu443 (TM7), as well as the polar contacts Asn675 and Lys679 (TM10–TM11), which together with the conserved tryptophan cage define the pocket geometry we hypothesized could contribute to isoform selectivity (see §2.7 for the full contact set (Martin et al., manuscript in preparation)).

Models corresponding to lumenal-facing occluded and cytosol-facing conformations were generated and refined to preserve key hydrogen-bonding networks and eliminate steric clashes. The resulting systems were solvated in explicit TIP3P water and subjected to molecular dynamics (MD) and Gaussian accelerated MD (GaMD) simulations in apo form and in complex with representative ligands (plosaracetam, LEV, and padsevonil).

Principal component analysis (PCA) of binding-site residues revealed that the SV2C pocket samples a relatively compact conformational space across apo and ligand-bound states. In contrast, SV2A exhibited greater conformational variability upon ligand binding (Figure 1). These observations suggest that the SV2C lumenal-facing occluded conformation may already resemble a binding-competent state for plosaracetam-like ligands, motivating its use as the primary receptor for virtual screening.

**Figure 1:**
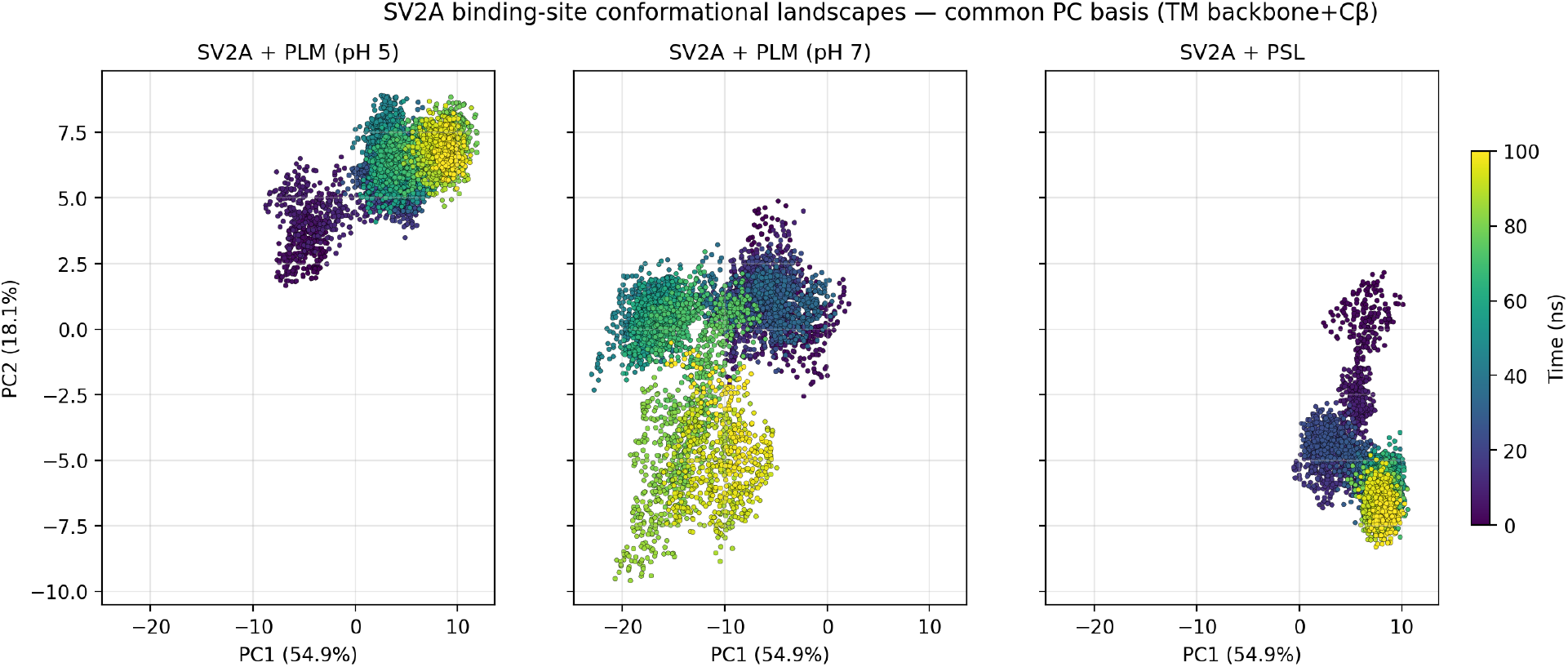
Principal Component Analysis (PCA) of SV2A conformational landscapes on a common PC basis derived from the concatenated three-trajectory ensemble (TM backbone+C*β* heavy atoms, 1,804 atoms). (a) Plosaracetam (PRM) at pH 5, (b) PRM at pH 7, and (c) padsevonil (PSL) at pH 7. PRM at pH 7 exhibits a 4.81 × greater PC1+PC2 variance spread and 65% higher per-residue RMSF than PSL (Table S5), indicating that PRM induces broader conformational sampling of the SV2A binding pocket. PSL produces a markedly more compact ensemble, in contrast to the broader conformational dynamics observed with PRM.

The 21 SV2C residues forming the bound-state contact shell, comprising the conserved tryptophan cage (Trp286/TM5, Trp440/TM7, Trp651/TM10), an aromatic shell (Phe263, Tyr447, Tyr448) lining the lumen-facing wall, and a polar contact ring (Asn644, Thr647, Asn675, Asp655, Lys679) on TM10–TM11; full set in §2.7, anchor the binding-competent geometry sampled by our MD trajectories. Per-residue RMSF analysis on the common-basis TM backbone+C*β* ensemble (Table S6) reveals that the elevated dynamics of SV2A in complex with plosaracetam at pH 7 are concentrated on the TM7 and TM10 residues flanking the conserved tryptophan cage (Tyr461/Tyr462 at TM7; Gly659, Ser662, Ile663 at TM10; SV2A numbering), not on peripheral pocket lining. By contrast, the SV2A + padsevonil ensemble shows uniformly low per-residue RMSF across the same set, and the SV2C trajectory shows the lowest amplitude motion overall (Figure 2). This residue-resolved pattern is consistent with the SV2C model being poised in a binding-competent configuration (Figure 3) and indicates that the differences in dynamics observed between plosaracetam and padsevonil are anchored at the conserved aromatic cage, the structural element with the highest pharmacological consequence.

**Figure 2:**
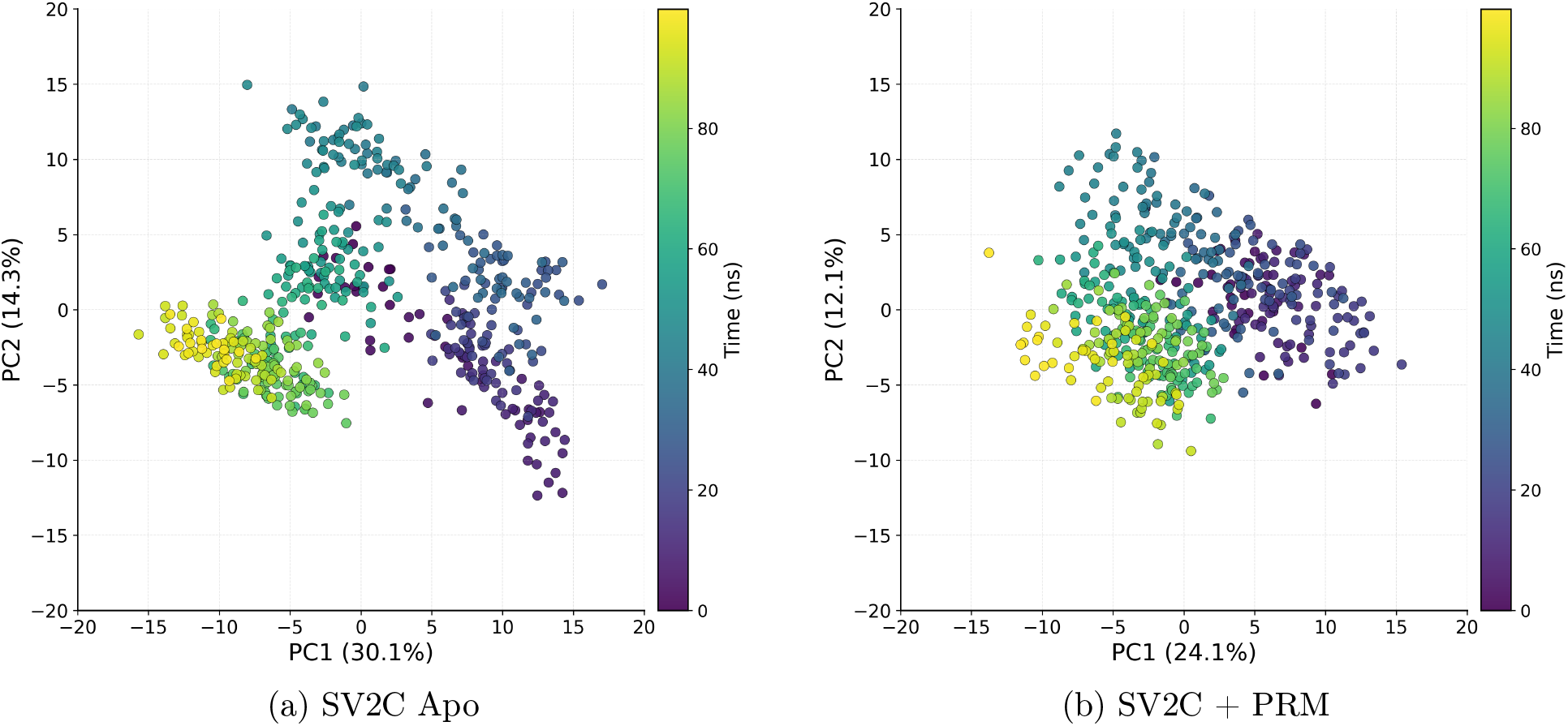
PCA conformational landscapes for the SV2C transmembrane bundle from 100 ns trajectories at pH 7. (a) In the apo state, SV2C exhibits a diffuse and disordered conformational ensemble. (b) Upon binding plosaracetam, the protein undergoes a dramatic structural ordering, transitioning to a highly compact and stable cluster. This contrast underscores plosaracetam’s role as a structural scaffold that stabilizes the binding-competent state of SV2C.

**Figure 3:**
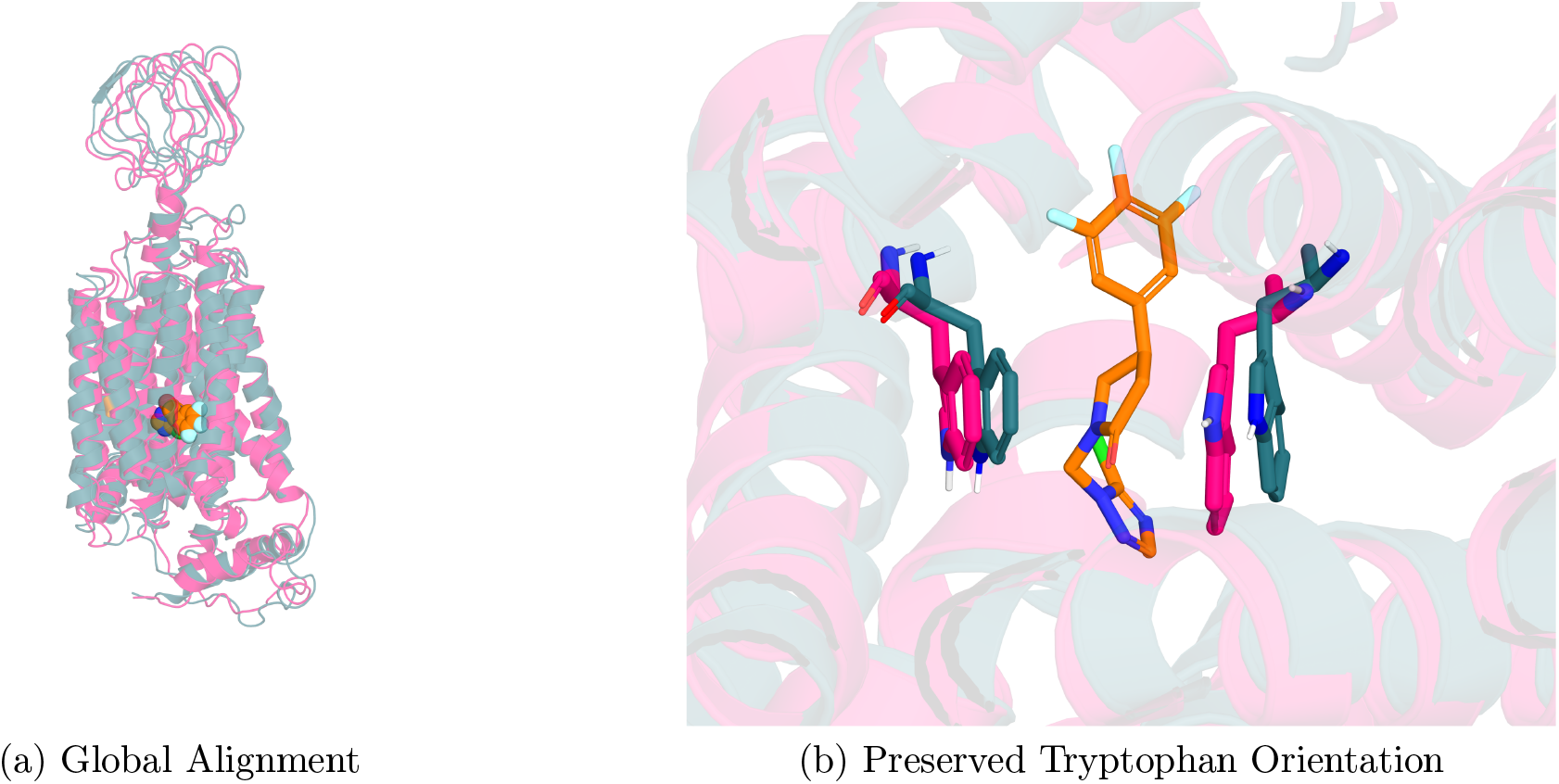
Structural comparison of SV2C in the apo state (magenta) and the plosaracetam-bound holo state (teal). (a) Global alignment of the two states demonstrates high structural resemblance, with plosaracetam (bright orange spheres) positioned within the transmembrane bundle. (b) A detailed view of the binding site reveals that the tryptophan residues maintain a nearly identical orientation in both states. This preserved geometry provides the binding-competent configuration required for stable coordination of the PRM trifluorophenyl group.

### 2.2 Retrospective Validation of the Virtual Screening Pipeline

A convolutional neural network-based scoring function (CNN_VS)[13], benchmarked retrospectively on a curated 39-ligand SV2A dataset (Methods), demonstrated strong predictive performance, yielding a Pearson correlation coefficient of *r* = 0.72 between predicted scores and experimental pIC_50_ values. This level of agreement provided confidence in the ability of the model to rank ligands effectively and justified its use as the primary scoring function in the prospective campaign, with free-energy methods reserved for refinement of top candidates.

### 2.3 Prospective Virtual Screening Campaign

Applying this validated pipeline to the Mcule in-stock library[14] (Methods) yielded a CNS-biased screening library of approximately 3.19 million compounds, which was ranked using a multi-stage, ligand- and structure-based funnel (Methods) to select 94 candidates for experimental testing (Figure 4).

**Figure 4:**
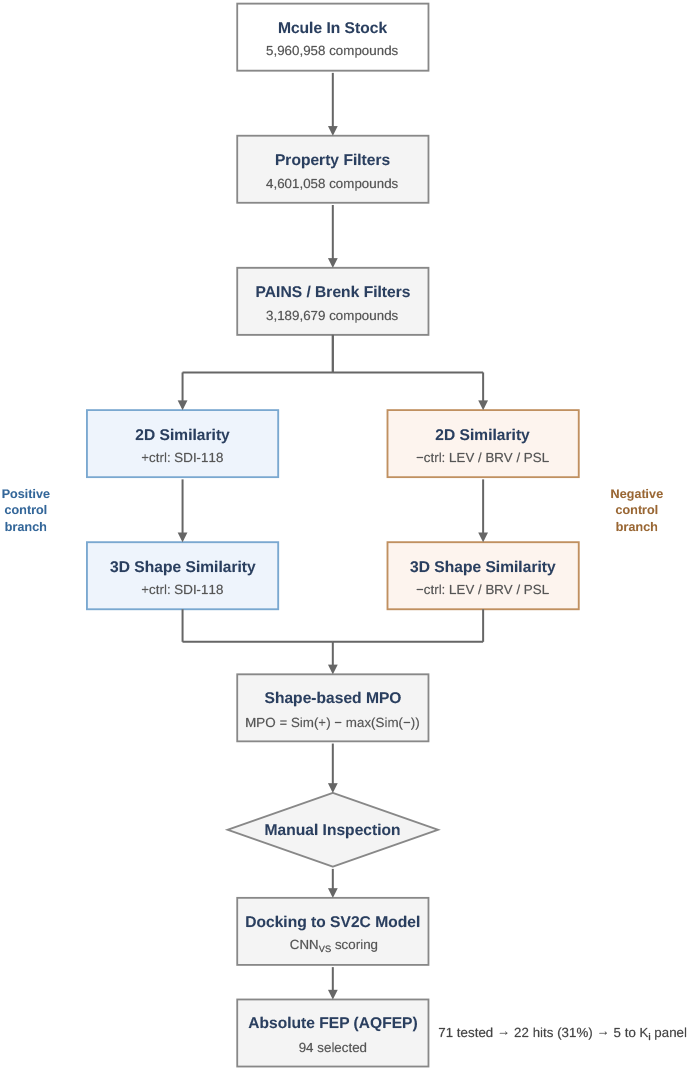
Multi-stage virtual screening funnel. Starting from the Mcule in-stock library (5,960,958 compounds), sequential property filters (aLogP, MW, HBA, HBD, stereocenters; 4,601,058 retained) and PAINS/Brenk alerts (3,189,679 retained) defined the CNS-biased screening library. Compounds were then ranked in parallel for 2D and 3D shape similarity to the positive control plosaracetam (PRM) and penalized for similarity to the negative controls LEV, BRV, and PSL; these scores were combined into a shape-based multi-parameter optimization (MPO) score [MPO = Sim(+ctrl) − max(Sim(−ctrls))]. After manual inspection and structure-based docking to the lumenal-facing occluded SV2C homology model (CNN_VS_ scoring), absolute binding free-energy calculations (AQFEP) refined the ranked list to 94 candidates. Of these, 71 were experimentally profiled, yielding 22 hits (31% hit rate), five of which advanced to isoform-selectivity *K*_*i*_ determination.

### 2.4 Biophysical Screening and Hit Identification

Of the 94 prioritized compounds, 71 were experimentally evaluated using an orthogonal assay cascade comprising a thermal shift assay (TSA) and a [^3^H]-padsevonil scintillation proximity assay (SPA). Each compound was tested at 100 *µ*M. Compounds were classified as primary hits if they displaced [^3^H]-padsevonil to a residual PSL *<*30% of the DMSO control and/or stabilized SV2C to ≥120% of the plosaracetam positive-control peak height in the thermal shift assay[15].

This campaign yielded 22 active compounds, corresponding to a 31% hit rate, indicating substantial enrichment relative to typical virtual screening efforts (Figure 5). Complete primary-cascade data for all 71 compounds, including inactive controls, are provided in the Supporting Information.

**Figure 5:**
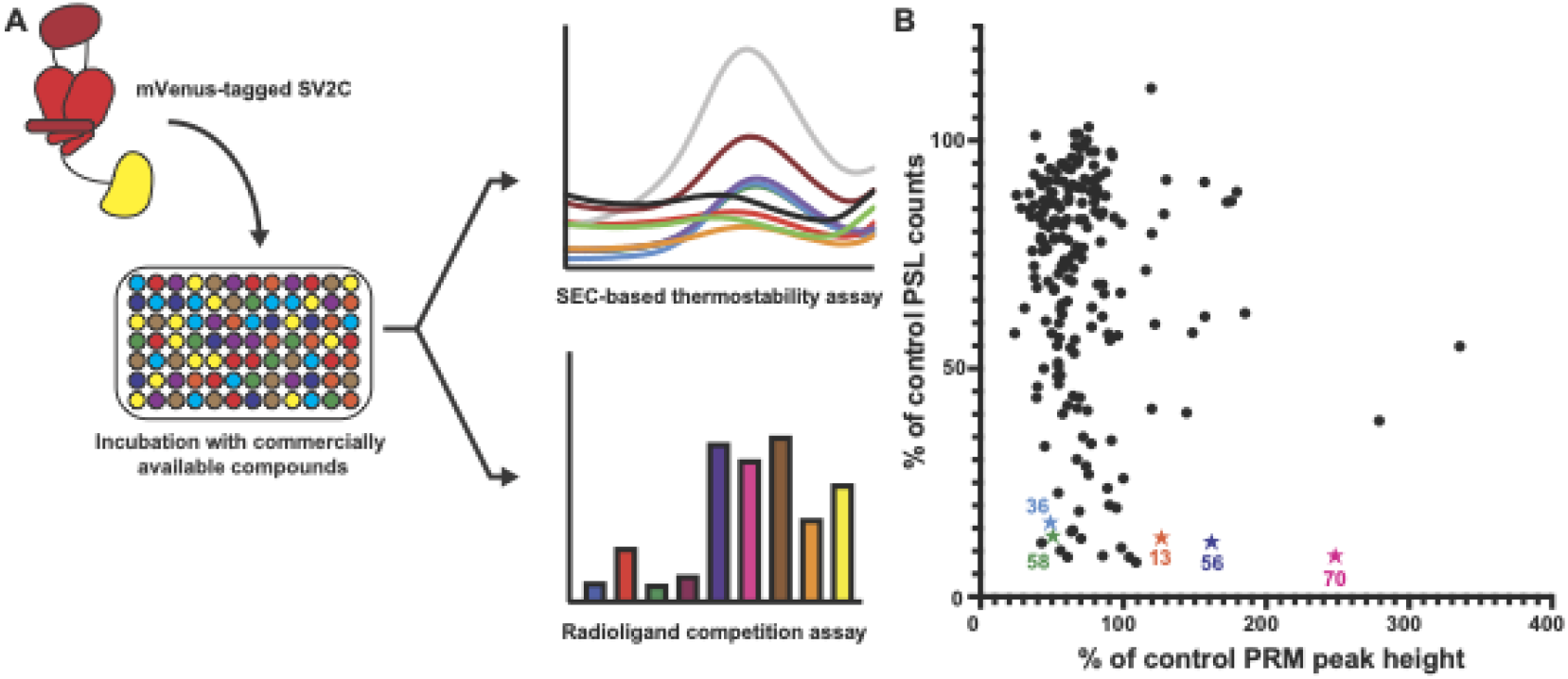
Binding of candidate ligands was examined by both thermostability and competition binding assays. (a) SV2C was heated in the presence of various ligands and compared with PRM or apo conditions. Thermostability was assessed using size exclusion chromatography, detecting a fluorescent mVenus tag on SV2C. Compounds that stabilize SV2C increase the peak height. Candidate compounds were also studied by competition binding with ^3^H-PSL. Compounds that bind to the same site as PSL result in lower % PSL binding. (b) Overall results of thermostability and competition binding screen, plotting percentage of control PRM peak height in a thermostability assay (x-axis) and percentage of PSL bound in a radioligand competition assay (y-axis). Several different groups of compounds were identified: compounds that compete for PSL binding and also thermostabilize SV2C; molecules that show a large degree of stabilization and intermediate PSL competition binding; and molecules that do not compete substantially but thermostabilize SV2C. Five compounds (13, 36, 56, 58, 70) which were advanced to isoform-selectivity profiling are highlighted.

### 2.5 Functional Classification of Hits

The identified hits segregated into functional classes based on their activity profiles in the two orthogonal assays:

#### *Competitors* (residual PSL *<*30%)

Compounds that robustly displaced [^3^H]-padsevonil from the orthosteric binding site. Thermostabilization activity (TSA ≥120% of the plosaracetam control) was observed in a subset of competitor compounds, confirming that padsevonil site occupancy and thermal stabilization are not tightly correlated assay readouts.

#### *Non-competitors* (residual PSL ≥30%)

Compounds that did not effectively displace [^3^H]-padsevonil from the orthosteric binding site, regardless of thermostability signal.

Padsevonil competition was used as the primary pharmacological discriminant; thermostabilization is reported as a secondary characterization without implying a specific mechanism.

Compounds in this thermostabilizing subset also qualitatively elevate steady-state SV2C protein levels in cells at 48 h (Martin et al., manuscript in preparation), suggesting that biophysical stabilization translates to a cellular pharmacochaperone-like effect.

### 2.6 Binding Affinity and Isoform Selectivity

Five first-screen hits that showed primary site competition and varied degrees of thermostabilization were advanced to radioligand competition binding assays against SV2A, SV2B, and SV2C using [^3^H]-padsevonil as the tracer. Inhibition constants (*K*_*i*_) were determined by one-site specific binding experiments in membranes enriched with SV2A, SV2B, or SV2C; the underlying dose-response curves are shown in Figure 6, and fitted values are summarized in Table 1 and Figure 7.

**Table 1:**
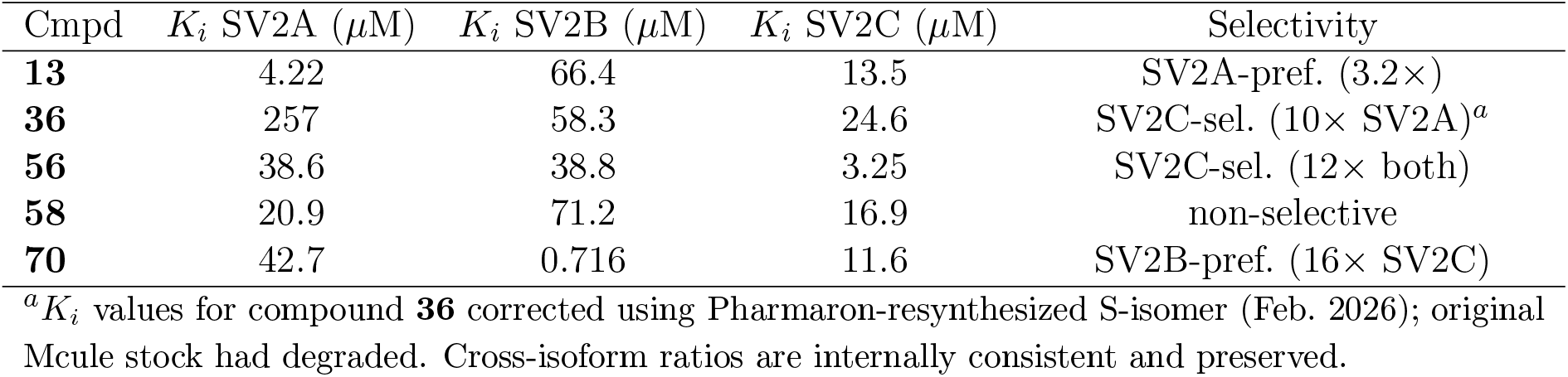
Binding affinities (*K*_*i*_) and isoform selectivity for first-screen leads determined by [^3^H]-padsevonil radioligand competition.

**Figure 6:**
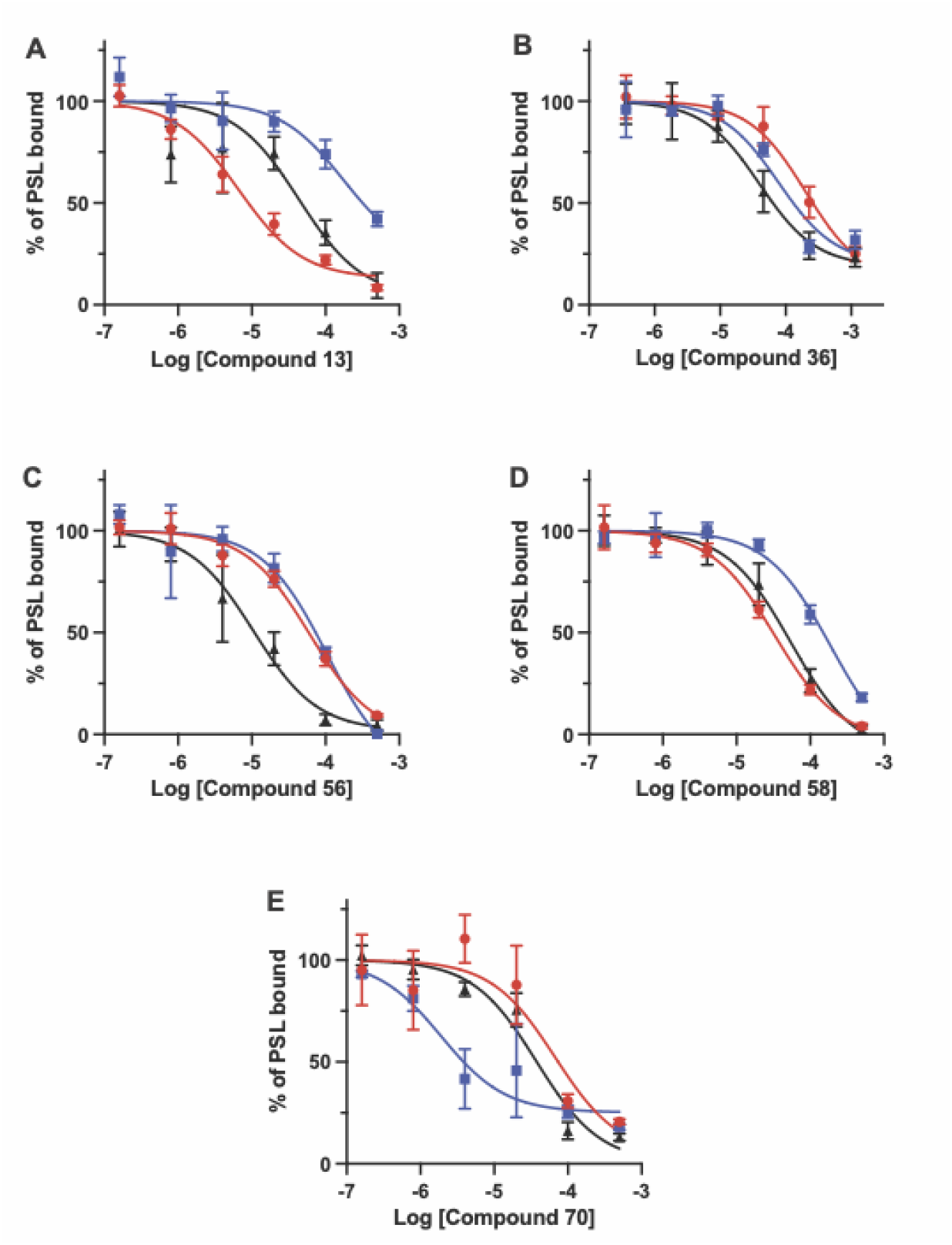
Competition binding of PSL with identified compounds with SV2A (red circles), SV2B (blue squares), or SV2C (black triangles) showing that compound 36 and 56 are selective to SV2C. Binding is plotted as a dose-response competition binding curve for compounds 13, 36, 56, 58, and 70, the *K*_*i*_ values summarized in Table 1.

**Figure 7:**
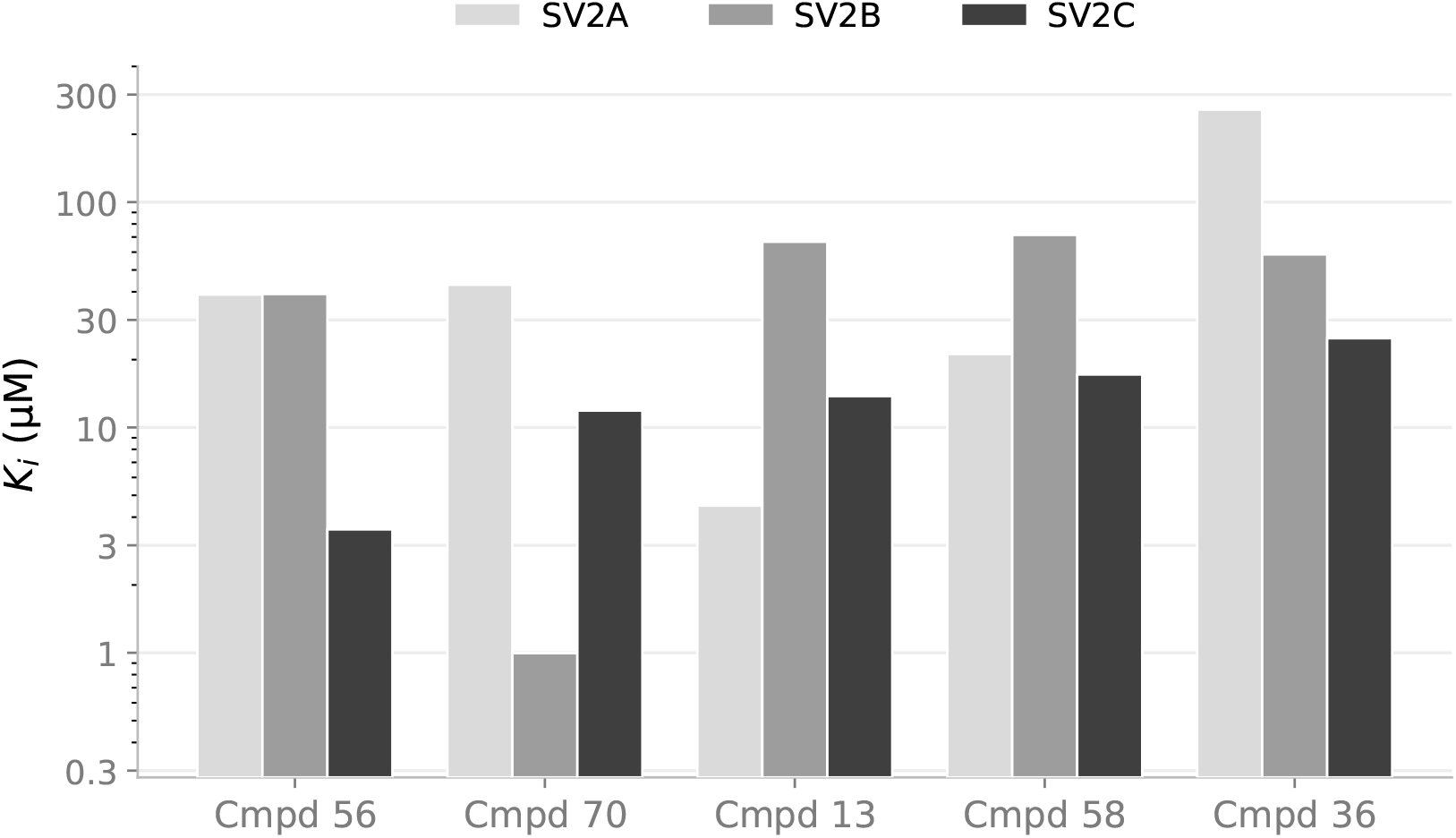
Isoform selectivity profile of five first-screen hits determined by [^3^H]-padsevonil radioligand competition binding. *K*_*i*_ values for SV2A (light grey), SV2B (mid grey), and SV2C (dark grey) are plotted on a logarithmic scale, sorted by SV2C affinity. Compounds **56** (*K*_*i*_ = 3.25 *µ*M, 12-fold selective over both SV2A and SV2B) and **36** (*K*_*i*_ = 24.6 *µ*M, 10-fold selective over SV2A) are the primary SV2C-selective leads. Compound **70** is an outlier with sub-micromolar SV2B affinity (*K*_*i*_= 716 nM; 16-fold selective over SV2C). Compounds **13** and **58** show modest SV2A preference and no meaningful isoform discrimination, respectively.

Compound **56** exhibited the best SV2C selectivity, with a *K*_*i*_ of 3.25 *µ*M for SV2C and approximately 12-fold selectivity over both SV2A and SV2B, making it the most promising lead for SV2C-directed optimization. Compound **36** showed a similar SV2C preference (24.6 *µ*M; ∼10-fold over SV2A) though with weaker discrimination against SV2B (∼2.4-fold). In contrast, compound **70** emerged as an unexpected SV2B-selective ligand, with a sub-micromolar *K*_*i*_ of 716 nM for SV2B, the strongest binding observed in either screen, and 16-fold selectivity over SV2C. The remaining compounds showed either modest SV2A preference (**13**) or no meaningful selectivity (**58**). Together, these data demonstrate that structurally distinct chemotypes accessed by the VS campaign can achieve divergent isoform selectivity profiles across the SV2 family (Figure 8), and they identify compounds **56** and **36** as the primary SV2C-selective leads entering hit-to-lead optimization.

**Figure 8:**
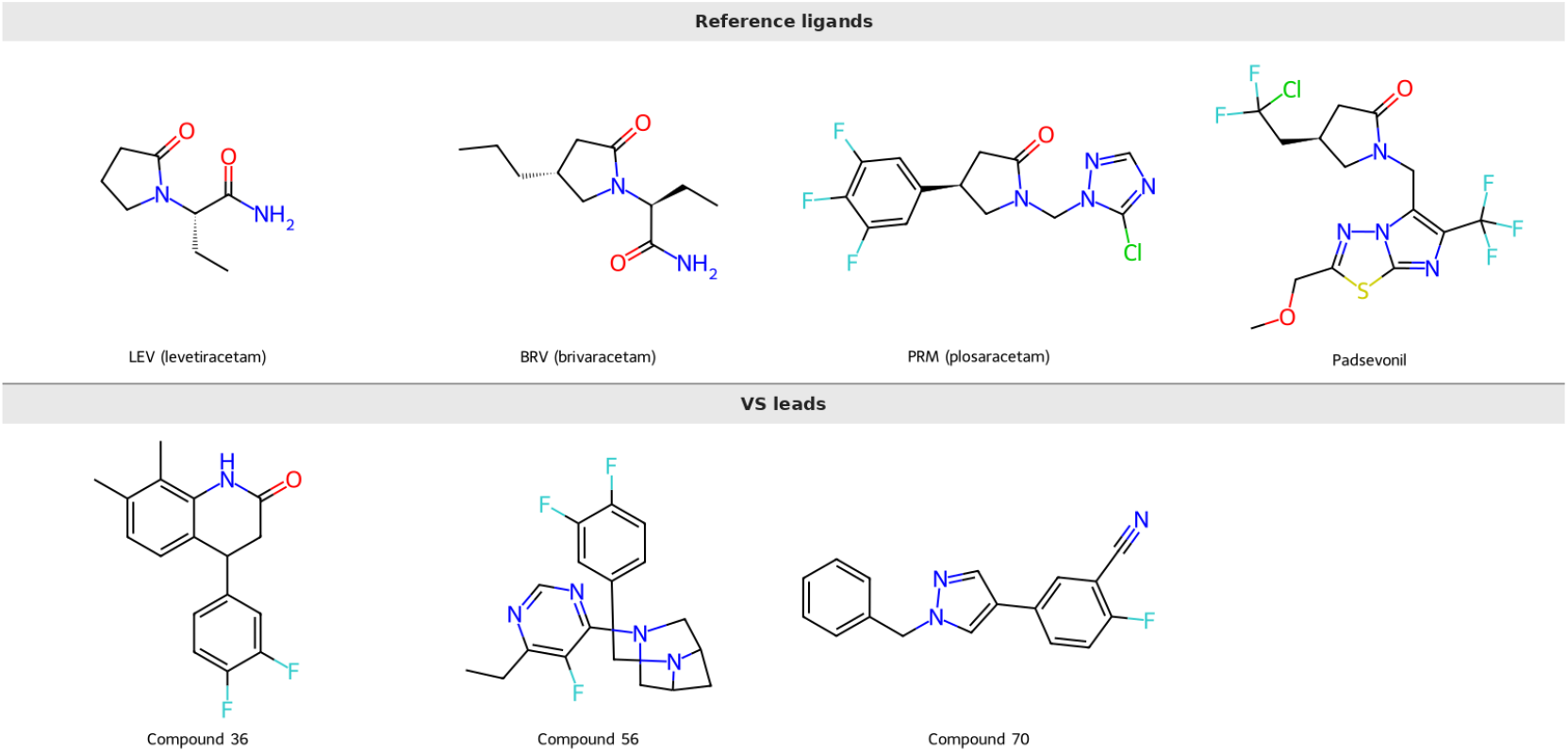
2D structures of three representative first-screen leads alongside established SV2-targeting reference ligands. *Top row* : reference compounds shown for structural context, including the SV2A-selective clinical racetams levetiracetam (LEV), brivaracetam (BRV), and plosaracetam (PRM), and the pan-SV2 ligand padsevonil (radiotracer used in this work). *Bottom row* : VS leads. Compounds **36** (SV2C-selective, 10×over SV2A) and **56** (SV2C-selective, 12× over both SV2A and SV2B) are the primary SV2C-selective leads. Compound **70** (SV2B-preferring, *K*_*i*_ = 716 nM) illustrates that closely related screening outputs can switch isoform preference. All three VS scaffolds feature fluorinated aromatic groups positioned to interact with the conserved tryptophan cage. Structures of the remaining two Ki-panel compounds (**13, 58**) are provided in the Supporting Information.

Notably, compound **70** showed strong competition with SV2C and an increase in thermostability, (§2.5), yet it is most selective for SV2B, binding at sub-micromolar affinity with 60-fold selectivity over SV2A and 16-fold selectivity over SV2C. This dissociation between functional class and isoform preference illustrates that the primary cascade, which only targeted SV2C, does not predict isoform selectivity. Compound **70** was resynthesized and initial results were confirmed in a retest (100 *µ*M, *>*98% [^3^H]-padsevonil competition against SV2C), ruling out batch-specific artifacts and establishing compound **70** as a validated SV2B-selective chemical tool.

No selective small-molecule SV2B ligands have been reported in the published literature to date; the existing pharmacology of SV2B is limited to observations with pan-SV2 ligands or with SV2A- and SV2C-preferring radioligands developed primarily for imaging applications (e.g. [^11^C]-UCB-J for SV2A and UCB-F for SV2C[9, 10]); none of these are SV2B-selective. SV2B is enriched in glutamatergic and GABAergic neurons, where it contributes to excitatory and inhibitory neurotransmission, and SV2B knockout mice display a seizure-prone phenotype[16], yet the absence of selective chemical tools has precluded pharmacological dissection of SV2B-specific function. Compound **70**, with its sub-micromolar SV2B affinity (*K*_*i*_ = 716 nM) and 16-fold selectivity over SV2C, provides a starting point for such studies. That a selective SV2B ligand emerged serendipitously from an SV2C-directed VS campaign underscores the chemical tractability of the SV2 family and suggests that the structural differences among isoform binding pockets, while subtle, are sufficient to support selective ligand discovery across the family.

### 2.7 Structural Basis of Binding and Selectivity

Docking analysis points to a shared binding orientation stabilized by conserved tryptophan residues within the SV2 pocket. These observations are supported by an unpublished cryo-EM structure of SV2A bound to plosaracetam (Martin et al., manuscript in preparation), which shows a binding-site C*α* RMSD of 0.76 Å relative to the SV2C homology model and a highly similar binding mode of the tryptophan residues. The 21-residue bound-state contact shell was identified from the PSL-SV2C structure (Martin et al., manuscript in preparation) using a 5 Å distance criterion around the bound ligand; the 20 highest-plasticity residues, with their SV2A cross-isoform equivalents, are provided in Table S6.

Sequence variations near the binding pocket, including SV2C Leu443 (TM7) and the polar contacts Asn675 and Lys679 (TM10–TM11), provide plausible structural determinants for selectivity. Additionally, per-residue RMSF analysis of the binding-site contact shell reveals that SV2A residues flanking the conserved tryptophan cage show substantially elevated dynamics in complex with plosaracetam relative to padsevonil — in particular Val608 (TM8; 1.35 Å PRM-pH7 RMSF), Tyr461/Tyr462 (TM7; 1.17–1.22 Å), and Gly659/Ser662/Ile663 (TM10; 0.86–1.03 Å) — whereas the corresponding SV2C ensemble is uniformly compact across the same residues (SV2A numbering; Table S6). This residue-resolved plasticity differential may impose stricter geometric constraints on ligand binding in SV2C and contribute to isoform discrimination.

### 2.8 Comparison to Prior SV2 Efforts and Implications

While SV2A has been extensively studied and pharmacologically exploited, SV2C has remained comparatively underexplored due to the lack of structural data and selective ligands. The present work demonstrates that an SV2A-derived structural framework, combined with dynamic modeling and AI-enhanced screening, can successfully identify SV2C-selective chemotypes.

The relatively high hit rate and emergence of multiple functional classes highlight the tractability of SV2C as a drug target and suggest opportunities for both orthosteric and allosteric modulation.

The cellular SV2C expression increase observed for the thermostabilizer subset (§2.5) extends the modulatory toolkit beyond direct binding and points to pharmacochaperone activity as an additional axis for SV2C-targeted chemistry.

### 2.9 Selective SV2C Inhibitors as a Complementary Therapeutic Axis

While the present campaign was framed around the discovery of selective SV2C positive modulators, the Competitors class identified in §2.5, compounds that robustly displaced [^3^H]-padsevonil with little or no thermostabilization of SV2C, represents a parallel and largely unexplored opportunity. These molecules are consistent with classical orthosteric site occupancy and were intentionally set aside as out of scope for the positive modulator brief. They were nevertheless characterized in the same primary cascade and, in several cases, advanced through *K*_*i*_ determination and isoform selectivity profiling against SV2A and SV2B. Treating these compounds as a complementary chemical-tool set, rather than as discarded byproducts, enables the same VS–experimental pipeline to address a distinct pharmacological hypothesis without additional discovery cost.

A selective SV2C inhibitor is mechanistically attractive in conditions characterized by elevated striatal dopamine signaling, including attention deficit hyperactivity disorder, Tourette syndrome, and other tic and movement disorders, as well as psychostimulant-related phenotypes. Because SV2C expression is sharply enriched in dopaminergic neurons of the basal ganglia, SV2C-selective therapies could modulate vesicular dopamine storage and release with greater precision than current vesicular monoamine transporter 2 (VMAT2) inhibitors used clinically for chorea and tardive dyskinesia. Further testing of these SV2C-specific compounds will provide insight into whether they function as inhibitors or through other mechanisms, and into the physiological effects of SV2C occupancy on dopaminergic neurotransmission. We anticipate that a small follow-on effort to assemble per-compound binding and selectivity tables for this subset, paired with functional readouts in dopamine-handling assays, will determine whether one or more chemotypes warrant dedicated hit-to-lead optimization in parallel with the modulator program.

### 2.10 Implications for Hit-to-Lead Optimization

The identified chemotypes represent structurally diverse starting points for medicinal chemistry optimization (Figure 8). Compound **56**, a piperazine-pyrimidine with 12-fold SV2C selectivity over both SV2A and SV2B, offers the most favorable selectivity profile and is the highest-priority scaffold for SAR exploration. Compound **36**, a dihydroisoquinolinone with 10-fold selectivity over SV2A but only 2.4-fold over SV2B, provides a complementary chemotype whose SV2B window may be widened through targeted modifications of the difluorophenyl substituent. Compound **70** constitutes the first reported selective small-molecule tool for probing SV2B biology and represents an independently valuable outcome of this campaign. Comparative SAR between compound **70** and the SV2C-selective leads may illuminate the structural determinants of isoform switching within closely related binding pockets and inform the rational design of isoform-selective ligands across the SV2 family.

Key optimization objectives include improving potency from the current low-micromolar range into the sub-micromolar regime while maintaining or improving isoform selectivity, and achieving CNS-relevant pharmacokinetic properties including brain penetration and metabolic stability. The observation that all three lead scaffolds feature fluorinated aromatic groups positioned to engage the conserved tryptophan cage suggests that modulating the electron density and geometry of these fluorinated moieties may provide a productive SAR axis for tuning both potency and selectivity.

More broadly, these compounds constitute the first generation of selective SV2C chemical tools for probing the role of this transporter in dopaminergic signaling and Parkinson’s disease biology. Given the limited biological characterization of SV2C function, further experiments are needed to determine what effect these molecules have on both SV2C function and neurotransmitter loading and release.

## 3 Conclusions

We have described the discovery of selective small-molecule binders for SV2C through an AI-enhanced virtual screening campaign integrated with medium-throughput biophysical characterization. Starting from an SV2A-derived homology model and a conformational ensemble generated by molecular dynamics and GaMD simulations, we screened 3.19 million CNS-biased compounds and experimentally validated 71 candidates, achieving a 31% primary hit rate. The resulting 22 active compounds classified into compounds that showed primary site competition with [^3^H]-padsevonil, and compounds that showed no primary site competition with padsevonil, demonstrating that structurally diverse SV2C ligands are accessible from a general commercial library. Radioligand competition binding against all three SV2 isoforms identified compounds **56** and **36** as SV2C-selective leads with 10–12-fold selectivity over SV2A, and compound **70** as a potent SV2B-selective ligand (*K*_*i*_= 716 nM). Principal component analysis of the SV2C binding site revealed a compact, binding-competent pocket geometry anchored by a conserved tryptophan cage (Trp286/TM5, Trp440/TM7, Trp651/TM10) and a polar contact ring (TM10–TM11). Per-residue RMSF analysis of the 21-residue binding-site contact shell identified substantially elevated dynamics in the SV2A site relative to SV2C, concentrated on residues flanking the tryptophan cage (Val608/TM8, Tyr461–462/TM7, Gly659–Ile663/TM10; SV2A numbering; Table S6), providing a residue-resolved structural rationale for isoform selectivity and a foundation for structure-guided optimization. Independent confirmation of the modeled binding site was provided by an unpublished SV2A–plosaracetam cryo-EM structure (Martin et al., manuscript in preparation), which revealed 19 of 21 identical contact residues and a binding-site C*α* RMSD of 0.76 Å relative to the SV2C homology model, validating both the structural framework and the fluorinated aromatic pharmacophore model that guided lead identification. These results establish SV2C as a tractable target for selective pharmacological intervention and deliver the first chemical tools for interrogating its role in dopaminergic signaling and Parkinson’s disease.

## 4 Methods

### 4.1 Homology Modeling and Multi-State Template Selection

Given the absence of high-resolution experimental structures for full-length synaptic vesicle glycoprotein 2C (SV2C), we developed a comprehensive structural framework using the high sequence identity (∼62%) and structural conservation within the SV2 family. We generated three-dimensional homology models of human SV2C (UniProt: Q496J9) representing the two primary functional orientations of the Major Facilitator Superfamily (MFS) transport cycle:

- **Lumenal-Facing Occluded State:** This model was generated using the human SV2A cryo-EM structure in complex with UCB-2500, a padsevonil analogue (PDB ID: 8UO9)[6], as the primary structural template; this structure represents a lumenal-facing occluded conformation. To account for the plasticity of the racetam binding site (RBS), an additional reference model was constructed using the brivaracetam-bound SV2A cryo-EM structure (PDB ID: 8K77)[12], which captures a more open lumen-facing state. This model was targeted to characterize the binding-competent orientation for luminal-facing modulators and served as the primary receptor for virtual screening.
- **Cytosol-Facing State:** To explore the alternative conformational extreme and evaluate activation-related structural shifts, we employed a hybrid-modeling strategy. The SV2A transmembrane (TM) core was threaded onto the crystal structure of the *E. coli* glycerol-3-phosphate transporter (glpT, PDB ID: 1PW4)[17], which provided a high-fidelity scaffold for the cytosol-facing helical arrangement.

Sequence alignment was performed, and initial 3D models were generated and refined using the *Modeller* suite[18]. We specifically prioritized the refinement of the racetam binding site, ensuring the optimized orientation of key binding-site residues (Trp300, Trp666, and Ile663 in SV2A; Trp286, Trp651, and Ile648 in SV2C) that define the hydrophobic core of the pocket. Loop regions, particularly the large luminal domain between TM6 and TM7, were energy-minimized to resolve steric clashes and optimize local hydrogen-bonding networks.

### 4.2 Molecular Dynamics (MD) Simulation Protocol

To transition from static snapshots to a dynamic conformational ensemble, we conducted all-atom molecular dynamics (MD) simulations using the OpenMM 8.0 engine[19]. Simulations were executed using an automated complex simulation workflow to ensure methodological reproducibility and standardized system preparation.

#### Force Field and System Setup

The protein systems were parameterized using the Amber ff14SB force field[20], and small-molecule ligands (including the reference ligand plosaracetam and prospective discovery hits) were parameterized with the General Amber Force Field (GAFF)[21]. The systems were solvated in a cubic box of TIP3P water[22] with a 10 Å buffer and neutralized with 0.15 M NaCl. To focus exclusively on the intrinsic protein-ligand stability and the conformational fluctuations of the TM bundle and luminal gate without the confounding influence of lipid-bilayer constraints, the simulations were conducted in a purely aqueous solvent environment.

#### Simulation Parameters

Each system underwent initial energy minimization using the L-BFGS algorithm, followed by a multi-step equilibration in the NVT and NPT ensembles at 310 K and 1 atm. Production MD runs were conducted for 100 ns with a 2 fs timestep. Bonds involving hydrogen atoms were constrained using the SHAKE algorithm, and long-range electrostatic interactions were handled via the Particle Mesh Ewald (PME) method[23] with a 9 Å cutoff. This duration was sufficient to observe structural convergence and the local ordering effects induced within the binding cavity upon ligand occupancy.

### 4.3 Principal Component Analysis (PCA)

To quantify the impact of ligand binding on the global conformational ensemble of SV2C, we performed Principal Component Analysis (PCA) on the Cartesian coordinates of the transmembrane backbone and C*β* heavy atoms (1,804 atoms) across the 100 ns trajectories, using a common PC basis fit to the concatenated three-trajectory ensemble to enable direct cross-condition comparison of conformational spread. This analysis allowed us to map the intrinsic breathing modes of the protein and assess the degree of structural ordering by projecting the trajectories onto the first two principal components.

### 4.4 Retrospective Benchmark Validation

Prior to prospective screening, we evaluated the computational workflow on a curated SV2A benchmark comprising 39 ligands with experimentally determined pIC_50_ values. Ligand poses were generated using maximum common substructure (MCS) alignment to an SV2A–BRV complex, followed by local minimization.

### 4.5 Virtual Screening Library Preparation

The Mcule in-stock library[14] (5.96 million compounds) was filtered in two sequential steps to enrich for CNS-relevant chemical space and remove undesirable chemotypes. First, physicochemical property windows consistent with CNS drug-like space were applied (aLogP 1–5, molecular weight ≤450 Da, hydrogen bond acceptors ≤10, hydrogen bond donors ≤5, and unspecified stereocenters ≤3), reducing the library to 4.60 million compounds. PAINS[24] and Brenk structural alerts were then applied, yielding a final screening library of approximately 3.19 million compounds.

### 4.6 Multi-Stage Virtual Screening Funnel

The screening workflow combined ligand-based and structure-based approaches in a multi-stage funnel. First, similarity and shape-based metrics were computed relative to plosaracetam, while penalizing similarity to SV2A-selective ligands (LEV, BRV) and the pan-SV2 ligand padsevonil; this comparison used 2048-bit Extended Connectivity Fingerprints (ECFP4)[25] to prioritize molecules mimicking the structural complementarity observed in the plosaracetam-bound SV2C trajectories, using the stabilized, ligand-bound MD conformation described above as the target receptor. This multi-parameter optimization enriched for chemotypes hypothesized to favor SV2C-like binding modes.

Top-ranked compounds were subsequently docked into the lumenal-facing occluded SV2C model, and poses were rescored using CNN_VS. A subset of candidates was further evaluated using free-energy methods to eliminate energetically unfavorable complexes. Final compound selection incorporated expert curation to ensure chemical diversity and tractability, yielding 94 compounds for experimental testing.

### 4.7 Cell Culture

tSA201 cells were maintained in suspension in SFM4Transfx-293 media (Cytiva) supplemented with 1% fetal bovine serum (Sigma) and 4 mM L-glutamine (Sigma). For adherent transfections, tSA201 cells were diluted into Dulbecco’s Modified Eagle Medium (DMEM) supplemented with 10% fetal bovine serum and 1 mM sodium pyruvate (Sigma). All cells were grown at 37 °C with 5% CO_2_.

### 4.8 Transient Transfections

For binding experiments using solubilized protein, adherent cells were transiently transfected according to previous methods[6, 26]. Briefly, pEG-BacMam plasmid containing full-length human SV2C linked to an 8×His-TwinStrep-mVenus tag was transfected into adherent tSA201 cells using PolyJet (SignaGen) following the manufacturer’s instructions. 48 hours post-transfection, cells were harvested, washed once with TBS150 buffer (20 mM Tris-HCl, pH 8.0, 150 mM NaCl), and pelleted. Pellets were stored at −80 °C for future use.

### 4.9 Production of SV2-Enriched Membranes

For membrane filtration assays, SV2A, SV2B, or SV2C were expressed according to previously published methods[6, 26]. Baculovirus with a titer of *>*10^9^ infectious particles/mL harboring the coding sequence for SV2AΔ64, full-length SV2B, or full-length SV2C, tagged with N-terminal (SV2B, SV2C) or C-terminal (SV2AΔ64) 8×His-TwinStrep-mVenus, was used to transduce suspension tSA201 cells. To enhance expression, sodium butyrate was added to a concentration of 10 mM 18–22 hours post-transduction. Cells were harvested 72 hours post-transduction and washed in TBS150, and cell pellets were stored at −80 °C. To produce cell membranes enriched with SV2A, SV2B, or SV2C, thawed cell pellets were resuspended in TBS150 and sonicated at 4 °C using a Fisherbrand Q500 sonicator (2 s on, 4 s off, 40% power, 15 min total sonication time). The sonicated mixture was cleared of large debris by centrifugation for 20 min at 12,000×*g*, followed by ultracentrifugation at 180,000×*g* for 2 hours to isolate membranes. Membranes were resuspended in TBS150 and homogenized by 25–30 passes through a Dounce homogenizer, then stored at −80 °C for future use.

### 4.10 Fluorescence Detection Size Exclusion Chromatography

All fluorescence detection size exclusion chromatography (FSEC)[15] was performed using a Shimadzu HPLC instrument with a refrigerated autosampler and RF20-AXS fluorescence detector, attached to a Superose 6 5/150 column (Cytiva). mVenus fluorescence was monitored in single-wavelength mode (515 nm excitation, 528 nm emission).

### 4.11 Scintillation Proximity Assays

For scintillation proximity assays, transiently transfected SV2C pellets were resuspended in TBS150 buffer containing 10 mM lauryl maltose neopentyl glycol (Anatrace, MNG) and 1 mM cholesteryl hemisuccinate (Anatrace, CHS). Pellets were solubilized for 90 min at 4 °C with mixing, and the solubilized mixture was cleared by ultracentrifugation for 30 min at 100,000×*g*. Fluorescently tagged SV2C in the whole-cell lysate was quantified using FSEC monitoring mVenus fluorescence. Lysates were diluted to a concentration of 5–10 nM SV2C in TBS150 buffer containing 0.1 mM MNG and incubated with 30–40 nM [^3^H]-padsevonil (specific activity 72 Ci/mmol, Pharmaron), 0.5 mg/mL Cu-YSi scintillant beads (Revvity), and 100 *µ*M of test compound or an equivalent volume of DMSO, for 2 hours or until control [^3^H]-PSL counts were stable. Non-specific counts were determined by addition of 100 *µ*M unlabelled padsevonil. All samples were measured in triplicate in 96-well plates using a MicroBeta 2 scintillation counter, and data were analyzed in GraphPad Prism 10.6.1[27].

### 4.12 Thermostability Assays

For thermostability assays, transiently transfected SV2C pellets were solubilized following the above procedure. 100 *µ*L samples of SV2C were incubated on ice for 30 min with 100 *µ*M of test compound or an equivalent volume of DMSO. After incubation, samples were heated in a PCR machine for 15 min. Heated samples were cleared with 96-well filtration plates (PALL Corporation) and analyzed by FSEC monitoring mVenus fluorescence. Percent of control peak height was determined by comparing the mVenus fluorescence intensity of DMSO-treated samples to samples treated with test compounds.

### 4.13 Membrane Filter Binding Assays

To determine isoform selectivity of candidate compounds, membranes enriched with SV2A, SV2B, or SV2C were incubated for 1 hour on ice in TBS150 containing 5 nM [^3^H]-padsevonil (specific activity 72 Ci/mmol) and serially diluted candidate compounds. 100 *µ*L of 2.5% polyethyleneimine (PEI, Sigma) solution in TBS150 was passed through each filter well of a glass fiber C 96-well filter plate (Revvity) using a vacuum manifold. After incubation, membranes were passed through the PEI-treated filters and washed four times with 100 *µ*L of TBS150. Non-specific background counts were determined by preparing samples containing membranes with 5 nM [^3^H]-PSL and 100 *µ*M unlabelled padsevonil. After washing, the plates were dried overnight. The next morning, 30 *µ*L of scintillant cocktail (MicroScint-PS, Revvity) was added to each filter and the plate was counted in a MicroBeta 2 scintillation counter. Inhibition constants (*K*_*i*_) were derived from one-site total binding fits using GraphPad Prism 10.6.1[27]. Each curve comprised triplicate determinations at each concentration. Goodness of fit was assessed by the coefficient of determination (*R*^2^); all reported *K*_*i*_ values correspond to fits with *R*^2^ ≥ 0.70.

## Abbreviations

SV2A/B/C: synaptic vesicle glycoprotein 2A/B/C
PD: Parkinson’s disease
VS: virtual screening
MD: molecular dynamics
GaMD: Gaussian accelerated molecular dynamics
TSA: thermal shift assay
SPA: scintillation proximity assay
CNS: central nervous system
PCA: principal component analysis
MFS: major facilitator superfamily
LEV: levetiracetam
BRV: brivaracetam
PRM: plosaracetam
PSL: Padsevonil
CNN: convolutional neural network
SAR: structure–activity relationship
VMAT2: vesicular monoamine transporter 2

## Acknowledgements

The authors thank Opher Kornfeld, Jeyanthi Ramasubbu, Monet Jimenez, and Ben Kraemer (SPARK NS) for strategic guidance and program management throughout this project. Dr. Miller and Dr. Bucher were also supported by NIH ES023839. This work was supported by the SPARK NS Translational Research Program.

## Author contributions

A.C.B. and M.F.M. contributed equally and designed the study. A.C.B. performed virtual screening, molecular dynamics simulations, and homology modeling, and wrote the manuscript. M.F.M. performed and analyzed the primary biophysical screening cascade and binding affinity assays. S.K. performed additional molecular dynamics analyses. B.S. performed ligand-based virtual screening. A.M. contributed to assay development and screening data. J.A.S. contributed to project management. R.S.F. and A.B. provided computational oversight and scientific direction. A.S. provided scientific direction for functional assays. M.L.B. contributed protein expression data. J.A.C. and G.W.M. supervised the study and obtained funding. All authors reviewed and approved the manuscript.

## Data availability

The virtual screening library (Mcule in-stock compounds) is available from Mcule Inc. Structural coordinates for the SV2C homology model, MD trajectories, and biophysical assay data supporting the findings of this study are available from the corresponding authors upon reasonable request.

## Code availability

Virtual screening, docking, and molecular dynamics analyses used SandboxAQ’s proprietary internal software, including the AQFEP method, which is not publicly available. Custom analysis scripts used to generate figures are available from the corresponding authors upon reasonable request.

## Supplementary Information

Compound identification table (MCULE catalog IDs, molecular formulae, SMILES, and primary-cascade functional classifications for all *K*_*i*_-panel leads), 2D structures for compounds **13** and **58**, full one-site total binding curve-fit parameters (log*K*_*i*_, SEM, *R*^2^) for all five first-screen leads against SV2A, SV2B, and SV2C, complete primary-cascade assay data ([^3^H]-padsevonil SPA and thermal shift) with SMILES for all 71 first-screen compounds including inactive controls, and SV2A conformational compactness metrics across common-basis PCA trajectories (Table S5).

