## Supplemental Information for "Discovery of Selective Small-Molecule Ligands of SV2C by AI-Enhanced Virtual Screening and Experimental Validation"

‖*These authors contributed equally.*


#### Compound Identification

Table S1 lists the five first-screen leads advanced to isoform-selectivity $K_i$ determination, with compound numbers as used in the main text, commercial catalog identifiers, molecular formulae, molecular weights, and primary-cascade functional classifications.

Table S1: Compound identification for first-screen $K_i$-panel leads.

| Cmpd | MCULE ID | Formula | MW (Da) | Class | SMILES |
|---|---|---|---|---|---|
| **13** | 3453066484 | $C_{16}H_{17}ClFN_3O$ | 321.8 | Dual-action | see below |
| **36** | 2544086857 | $C_{17}H_{15}F_2NO$ | 287.3 | Competitor | see below |
| **56** | 9160992168 | $C_{18}H_{19}F_3N_4$ | 348.4 | Competitor | see below |
| **58** | 2167134489 | $C_{16}H_{13}ClF_3N_3O$ | 355.7 | Competitor | see below |
| **70** | 5211192032 | $C_{17}H_{12}FN_3$ | 277.3 | Dual-action | see below |

**SMILES:**

**13**: `C1(=NC=CN1C1CN(C(=O)C1)CCC)C1=CC=C(C=C1Cl)F`

**36**: `C12C(C)=C(C)C=CC=1C(CC(=O)N2)C1C=CC(=C(F)C=1)F`

**56**: `N1(CC2CC(N2CC2C=CC(=C(F)C=2)F)C1)C1=NC=NC(CC)=C1F`

**58**: `N1(CCN(C(=O)C1)CC1C=CC(=CC=1F)F)C1=C(F)C=C(C=N1)Cl`

**70**: `C1(=CN(CC2C=CC=CC=2)N=C1)C1C=C(C#N)C(F)=CC=1`

### 2D Structures of All Ki-Panel Leads

Figure S1 shows 2D structures for all five compounds advanced to isoform-selectivity profiling. Compounds **36**, **56**, and **70** are shown in the main text (Figure 6); compounds **13** and **58** are shown here for completeness.

### Radioligand Competition Curve-Fit Parameters

Table S2 reports the full one-site total binding fit parameters from [$^3$H]-Padsevonil radioligand competition assays for the five first-screen leads against SV2A, SV2B, and SV2C. All fits were performed using GraphPad Prism. Each fit comprised 21–24 data points (triplicate determinations at 7–8 concentrations).

### Full First-Screen Assay Data

Table S3 reports the primary-cascade assay readouts for all 71 compounds evaluated in the first screen. Each compound was tested at 100 $\mu$M in both the [$^3$H]-Padsevonil scintillation proximity assay (SPA; % of [$^3$H]-PSL counts remaining relative to DMSO control) and the SV2C thermal shift assay (TSA; % of SV2C peak height after heating relative to the plosarac-

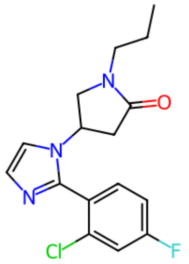

Compound 13

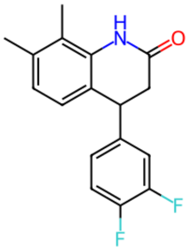

Compound 36

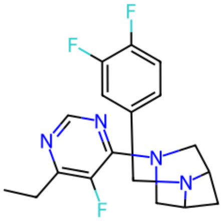

Compound 56

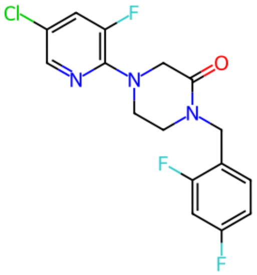

Compound 58

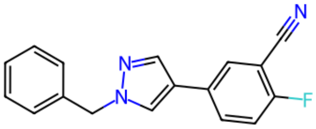

Compound 70

Figure S1: 2D structures of all five first-screen leads advanced to three-isoform $K_i$ determination.

Table S2: One-site total binding fit parameters for first-screen $K_i$-panel leads. $K_i$ values and log$K_i$ are best-fit values; SEM is the standard error of the log$K_i$ estimate; $R^2$ is the coefficient of determination. Compound **36** values reflect the Pharmaron-resynthesized S-isomer (Feb. 2026); the original Mcule stock had degraded, and the reported parameters incorporate the corresponding uniform concentration correction (log$K_i$ shifted, SEM and $R^2$ unchanged), consistent with Table 1 of the main text.

| Cmpd | Isoform | log$K_i$ | SEM | $K_i$ ($\mu$M) | $R^2$ |
|---|---|---|---|---|---|
| **13** | SV2A | −5.374 | 0.069 | 4.22 | 0.963 |
| **13** | SV2B | −4.178 | 0.194 | 66.4 | 0.820 |
| **13** | SV2C | −4.871 | 0.234 | 13.5 | 0.703 |
| **36** | SV2A | −3.590 | 0.091 | 257 | 0.942 |
| **36** | SV2B | −4.234 | 0.112 | 58.3 | 0.910 |
| **36** | SV2C | −4.609 | 0.099 | 24.6 | 0.926 |
| **56** | SV2A | −4.413 | 0.053 | 38.6 | 0.979 |
| **56** | SV2B | −4.411 | 0.108 | 38.8 | 0.923 |
| **56** | SV2C | −5.488 | 0.099 | 3.25 | 0.928 |
| **58** | SV2A | −4.679 | 0.050 | 20.9 | 0.981 |
| **58** | SV2B | −4.147 | 0.078 | 71.2 | 0.967 |
| **58** | SV2C | −4.773 | 0.063 | 16.9 | 0.971 |
| **70** | SV2A | −4.370 | 0.195 | 42.7 | 0.778 |
| **70** | SV2B | −6.145 | 0.174 | 0.716 | 0.810 |
| **70** | SV2C | −4.936 | 0.084 | 11.6 | 0.947 |

etam (PRM) positive control). SMILES strings for all compounds are listed in Table S4. All compounds were sourced from the Mcule in-stock library.

Table S3: Primary-cascade assay data for all 71 first-screen compounds. PSL% = percentage of [$^3$H]-Padsevonil counts remaining at 100 $\mu$M (lower values indicate stronger Padsevonil displacement). TSA% = percentage of SV2C peak height relative to the PRM control (values >100% indicate protein stabilization).

| Cmpd | MCULE ID | PSL (%) | TSA (%) |
|---|---|---|---|
| **1** | 1744011151 | 51.3 | 54.3 |
| **2** | 8486573340 | 74.2 | 70.9 |
| **3** | 9575564045 | 56.4 | 66.0 |
| **4** | 6491773015 | 88.8 | 48.9 |
| **5** | 4613988052 | 40.1 | 57.5 |
| **6** | 4223237215 | 26.8 | 75.7 |
| **7** | 9589926861 | 50.4 | 54.6 |
| **8** | 3723740449 | 69.7 | 61.3 |
| **9** | 3204374392 | 28.7 | 73.9 |
| **10** | 4682090687 | 33.5 | 77.5 |
| **11** | 7338887236 | 53.4 | 65.7 |
| **12** | 4770676454 | 40.8 | 74.8 |
| **13** | 3453066484 | 12.9 | 126.7 |
| **14** | 3062695716 | 76.8 | 66.9 |
| **15** | 6518104966 | 78.8 | 42.1 |
| **16** | 5836114029 | 86.5 | 172.5 |
| **17** | 8163149146 | 86.1 | 48.8 |
| **18** | 4987883590 | 93.8 | 48.6 |
| **19** | 3927732425 | 19.5 | 95.2 |
| **20** | 8358385252 | 59.2 | 77.8 |
| **21** | 3884805535 | 61.4 | 157.0 |

| Cmpd | MCULE ID | PSL (%) | TSA (%) |
| --- | --- | --- | --- |
| **22** | 2995652217 | 88.8 | 179.4 |
| **23** | 4636787591 | 72.5 | 37.6 |
| **24** | 1815640356 | 59.9 | 55.0 |
| **25** | 4234782027 | 43.6 | 69.7 |
| **26** | 4805543545 | 91.0 | 58.1 |
| **27** | 5074942139 | 83.9 | 128.5 |
| **28** | 2070751252 | 62.2 | 185.1 |
| **29** | 5063231354 | 67.8 | 39.6 |
| **30** | 5628969932 | 83.3 | 40.4 |
| **31** | 2657561688 | 84.1 | 43.7 |
| **32** | 5766304524 | 50.0 | 44.4 |
| **33** | 3617784916 | 38.6 | 279.1 |
| **34** | 9459092498 | 86.7 | 175.6 |
| **35** | 6466270040 | 43.6 | 39.3 |
| **36** | 2544086857 | 16.3 | 48.9 |
| **37** | 6690519334 | 54.9 | 335.5 |
| **38** | 9396775507 | 46.6 | 54.7 |
| **39** | 9272733791 | 57.9 | 148.4 |
| **40** | 8969059621 | 79.6 | 120.0 |
| **41** | 5824082902 | 11.8 | 42.8 |
| **42** | 9247407452 | 69.1 | 47.4 |
| **43** | 5974074158 | 91.4 | 42.3 |
| **44** | 3852793479 | 88.6 | 50.7 |
| **45** | 8462804602 | 96.1 | 69.2 |
| **46** | 2785659352 | 86.7 | 61.5 |
| **47** | 7347796145 | 63.1 | 56.1 |

| Cmpd | MCULE ID | PSL (%) | TSA (%) |
|---|---|---|---|
| **48** | 8120007357 | 76.0 | 67.7 |
| **49** | 2915246276 | 14.2 | 63.5 |
| **50** | 4321741950 | 10.1 | 56.0 |
| **51** | 2117441689 | 41.9 | 61.0 |
| **52** | 6436461392 | 34.3 | 91.6 |
| **53** | 8172552254 | 67.6 | 51.4 |
| **54** | 4534324501 | 76.2 | 46.7 |
| **55** | 1742432895 | 55.1 | 53.8 |
| **56** | 9160992168 | 9.0 | 85.7 |
| **57** | 5482316412 | 69.1 | 64.3 |
| **58** | 2167134489 | 13.3 | 50.7 |
| **59** | 7427365850 | 56.7 | 55.1 |
| **60** | 3831727223 | 35.0 | 71.9 |
| **61** | 5371251748 | 67.2 | 51.6 |
| **62** | 6200523234 | 82.8 | 60.6 |
| **63** | 8759114888 | 91.4 | 130.1 |
| **64** | 2892372674 | 78.7 | 51.2 |
| **65** | 5201651587 | 86.9 | 44.0 |
| **66** | 2172297089 | 84.3 | 85.7 |
| **67** | 4455050635 | 71.5 | 115.9 |
| **68** | 8732075272 | 94.0 | 60.4 |
| **69** | 1829398364 | 72.1 | 58.5 |
| **70** | 5211192032 | 9.0 | 248.2 |
| **71** | 7679901575 | 91.6 | 72.8 |

Table S4: SMILES strings for all 71 first-screen compounds (parent structures; counterions removed where applicable).

| Cmpd | SMILES |
|---|---|
| **1** | C1N(C(C2C=CC=C(C(F)(F)F)C=2C1)=O)CCOC1 |
| **2** | C1(C=NNC=1CC1CCCN(C(=O)C2=CC=CS2)C1)S(C)(=O)=O |
| **3** | C1(C(C1)C)C(=O)N1CCC(C1)C1=CC(O)=NC(N(C)C)=N1 |
| **4** | C1(NC(C)C)N=C(C=C(O)N=1)C1CC(N(CC2=CC=CC=N2)C1)=O |
| **5** | C1(=NC2C(=O)N(C3CCC(CC3)N(C)C)C=C2N=1)C1=CC=C(C=C1)C(F<br>)(F)F |
| **6** | C1(=NC(=O)C(F)(F)C2=CC=CC=C12)NCCN1CCCC(O)C1 |
| **7** | C1(C=CC(=CC=1)N1C(=O)NC2C(=NN=C12)C)OCC1=CN(C=N1)C1CCC<br>CC1 |
| **8** | C1(=CC(OC(=O)C2=CC(=CC=C2)F)=C2C(=O)OCC2=C1)F |
| **9** | C1(CCC(=O)N1CC1=CC=C(S1)S(=O)(=O)C)C1=CC2=CC=CC=C2C=C1 |
| **10** | C1(CC(=O)N2C(=O)C=CC2=C1)C1=CC=C(C=C1)F |
| **11** | C1(=NN=C(N1CC(=O)N1CCCC1)CC)C1=CC(=CC(=C1)Cl)Cl |
| **12** | C1(=NN=C(N1CC(=O)N1CCC(C1)O)CC)C1=C(Cl)C=CC=C1 |
| **13** | C1(=NC=CN1C1CN(C(=O)C1)CCC)C1=CC=C(C=C1Cl)F |
| **14** | C1(CN(CCC(N1)=O)C(=O)C1C=CC=CC=1)NCC1CCCO1 |
| **15** | C1(=NC=CN1C1CN(C(=O)C1)C)C1=C(Cl)C=CC(=C1)F |
| **16** | C1(CC(NS(=O)(=O)CC)CO1)CC(F)C1=CC=CC=C1 |
| **17** | C1(F)=C(C=CC(=C1)C1(O)CC1)NC1=NC=CC=C1F |
| **18** | C1(F)=CC=C(NC(=O)CC2=NNC(=O)C2)C(=C1)F |
| **19** | C1(OC(CC1CC1C=CC(=CC=1)F)C1=CC=C(C=C1)F)=O |
| **20** | C1(=CC(C(F)(F)F)=NN1CC1=CC(N)=NC=C1)C1CCCC1 |
| **21** | C1(CC(F)(F)F)(CC(OCC2=CC=CC=C2)C1)C(F)(F)F |
| **22** | C1(CC(=O)NCC)=NN(C=N1)C1CCSCC1 |

| Cmpd | SMILES |
|---|---|
| **23** | C1(=CC(=CC=C1F)F)C1=CC=C(S1)C#N |
| **24** | C1(=CC2=CC=CC=C2S1)NC(=O)CC1CCCC1 |
| **25** | C1(OC(CC1CC1C=CC(=CC=1)F)CC(F)(F)F)=O |
| **26** | C1(=CC=C(C=C1NC(=O)CC1CCOC1)C(F)(F)F)F |
| **27** | C1(=CC=C(C=C1)OCC1=NN(N=N1)C1CCCC1)C(N)=O |
| **28** | C1(=NC=C(N1CC1CCC(N(CC)CC)CC1)F)F |
| **29** | C1(=NC=CN1C1CCN(C(=O)C1)C)C1=C(Cl)C=CC(=C1)F |
| **30** | C1(=C(C)C=CC(=C1C)F)C(CC(=O)NCC1=NC=CC=C1)F |
| **31** | N1(C(=O)CNC(=O)C1CC1=CC=C(S1)C)CC1=NC=C(C=N1)N |
| **32** | C1(CC(=O)N2C(CCCS2(=O)=O)C1)C1=CC=CC(=C1)C |
| **33** | C1(OC(CC(O1)C1=CC=C(C=C1)F)COC(F)F)=O |
| **34** | C1(=CC=C(C=C1Cl)C1=NN2CCCCC2=N1)OC |
| **35** | C1(CC2=CC=CC=C2)OC(=O)C2=CC(=CC=C12)OC1=CC(=CC=C1)Cl |
| **36** | C12C(C)=C(C)C=CC=1C(CC(=O)N2)C1C=CC(=C(F)C=1)F |
| **37** | C1(=NC=CN1C1CCN(C(=O)C1)C)C1=C(Cl)C=C(C=C1)F |
| **38** | C1(=C(C)C=CC(=C1C)F)C(CC(=O)N1CCNCC1)F |
| **39** | C1(=C(C)C=CC(=C1C)F)C(CC(=O)N1CCN(C)CC1)F |
| **40** | C1(=C(C)C=CC(=C1C)F)C(CC(=O)NCC1=NC=CC=C1)F |
| **41** | C1(C(N(C2=CC=CC=C2)C=N1)CC1=CC=CC=C1)=O |
| **42** | C1(=CC=C(C(=O)S1)NCC1CCCCN1C)CC |
| **43** | C1(=CC(=C(O1)C)F)C(CC1=CC=CC=C1)=O |
| **44** | C1(=NC=C(N1CCC(NCC1CCCC1)=O)F)F |
| **45** | C1(=C(CCNC(=O)C2CCN(C(=O)C(C)(C)N)C2)NC=N1)OC |
| **46** | C1(=CC=C(S1)C1=CC(=CC(=C1)F)F)C(=O)NC1=NC2=CC=CC=C2N=C1 |

| Cmpd | SMILES |
|---|---|
| **47** | C1(=NC(=NC(C(NC2=CC=CC=C2)=O)=C1Cl)N1CCN(CC1)CC1=CC=CC=C1)F |
| **48** | C1(=CC2=CC=CC=C2S(=O)(=O)N1CC1CCCC1)CC(=O)NCC1=CC=CC=C1 |
| **49** | C1(=CC(=C(F)C=C1F)Cl)NC(=O)C1=CC=CS1 |
| **50** | C1(=NC=CC(=N1)Cl)N1CCN(CC1)CC1=CC=C(C=C1)F |
| **51** | C1(F)=CC=C(NC(=O)C2(CO)CN(C2)C(=O)OC)C(=C1)F |
| **52** | C1(=CN(N=C1CC1=CC=C(C=C1)F)C)C#N |
| **53** | C1(=CC(=C(Cl)C=C1)F)NC(=O)C1CC1 |
| **54** | C1(=CC=C(NC(=O)C2=CC(F)=CC(=C2)F)C=C1)OC |
| **55** | C1(=NN=C(N1CC(=O)N1CC(C1)O)CC)C1=C(Cl)C=CC=C1 |
| **56** | N1(CC2CC(N2CC2C=CC(=C(F)C=2)F)C1)C1=NC=NC(CC)=C1F |
| **57** | C1(CCCC(C1)OC(=O)C1=CC=CS1)OC1=CC=CC=C1 |
| **58** | N1(CCN(C(=O)C1)CC1C=CC(=CC=1F)F)C1=C(F)C=C(C=N1)Cl |
| **59** | C1(=NC=CN1C1CCN(C(=O)C1)CC1=CC(=CC=C1)F)C1=C(Cl)C=CC(=C1)F |
| **60** | C1(=NC=C(N1CCC(N1CCCC1)=O)F)C1=CC=C(C=C1)F |
| **61** | C1(OC(CC1CC1C=CC(=CC=1)F)CCC(F)F)=O |
| **62** | C1(CCNS(=O)(=O)CC(F)(F)F)NCC(N1)OC |
| **63** | C1(=CC(=CC(=C1)F)C1CC(=O)OC1)OC |
| **64** | C1(CC(N=C1)N1CCNC(=O)C1)C1=CC=CC(=N1)N |
| **65** | C1(=CC(=CC(=C1)F)C(F)(F)F)C#CC1=NC=CS1 |
| **66** | C1(CC(=O)NCC2C=CC(=C(OC)C=2)OC)N(C(=O)C2C=CC=CC=2)CC1=O |
| **67** | C1(=CC=C2N(C(CC2=C1)CC1=CC(=CC=C1OC)O)C=O)O |
| **68** | C1(=NN=C(N1CC(=O)NC1=CC(=CC=C1)F)C)C1=CC(=CC=C1)F |

| Cmpd | SMILES |
|---|---|
| **69** | `C1(NC2=CC=CC(=C2C1=O)F)C1=CC=CC(=C1)Cl` |
| **70** | `C1(=CN(CC2C=CC=CC=2)N=C1)C1C=C(C#N)C(F)=CC=1` |
| **71** | `C1(=CC=C(NC(=O)C2(CO)CC2)C(=C1)F)F` |

### SV2A Conformational Compactness Metrics

Table S5 reports quantitative compactness metrics for the three SV2A trajectories analyzed on a common PC basis (Figure 1 of the main text). All three metrics independently confirm that padsevonil (PSL) produces the most compact conformational ensemble.

Table S5: Quantitative compactness metrics for SV2A conformational ensembles on a common PC basis. Ratios are relative to PSL (most compact ensemble = 1.00×). Selection: TM backbone+C$\beta$ heavy atoms (1,804 atoms); common-basis PCA fit on concatenated 15,000 frames.

| Condition | RMSF (Å) | PC1+PC2 trace | Hull area | RMSF ratio | Trace ratio |
|---|---|---|---|---|---|
| PRM (pH 5) | $1.31 \pm 0.88$ | 17.30 | 96.9 | 1.18× | 2.29× |
| PRM (pH 7) | $1.84 \pm 1.42$ | 36.37 | 246.5 | 1.65× | 4.81× |
| PSL | $1.12 \pm 0.65$ | 7.56 | 84.5 | 1.00× | 1.00× |

### Cytosol-Facing vs Lumen-Facing Occluded Conformational Landscapes

Figure S2 compares the conformational landscapes of the SV2C cytosol-facing model (with plosaracetam, PRM) and the SV2A lumen-facing occluded trajectory (with padsevonil, PSL) on a common PC basis.

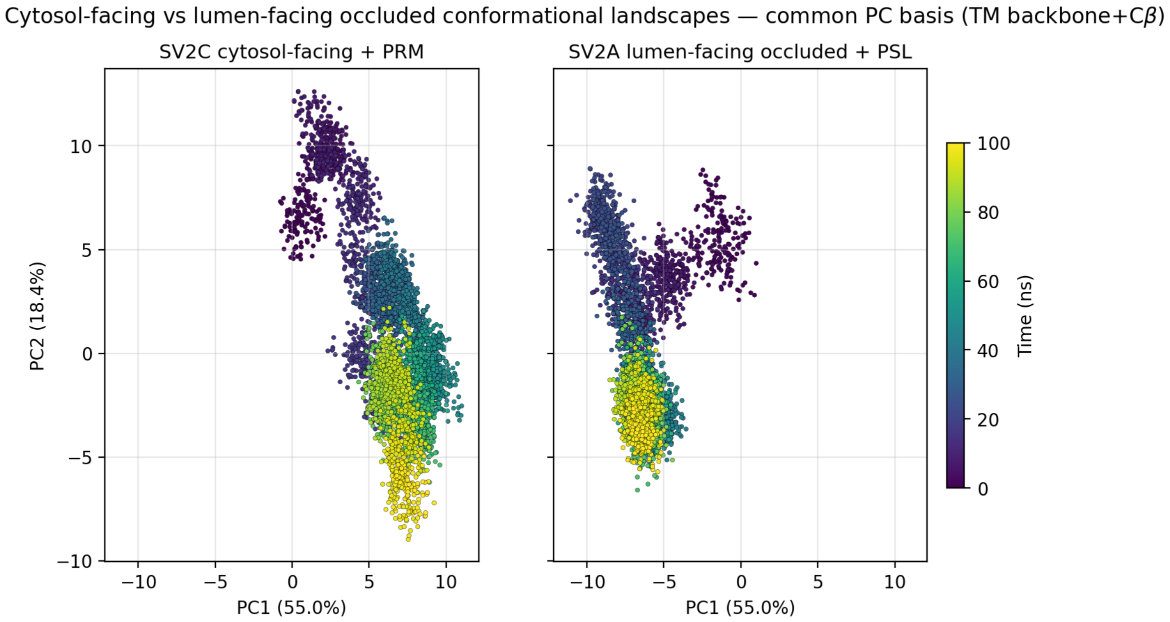


Figure S2: Conformational landscapes of the SV2C cytosol-facing model (with plosaracetam, PRM) and the SV2A lumen-facing occluded trajectory (with padsevonil, PSL) on a common PC basis (TM backbone+C$\beta$, 1,800 atoms; PC1+PC2 = 73.4% of variance). The cytosol-facing model samples a restricted region of positive PC1 throughout the 100 ns trajectory, while the lumen-facing occluded ensemble explores a broader conformational range including a distinct negative-PC1 basin in later frames. The partial overlap in early lumen-facing frames reflects the shared transmembrane core, whereas the late-frame divergence along PC1 corresponds to lumen-facing-specific dynamics not sampled by the cytosol-facing conformation.

### Per-Residue RMSF for Binding-Site Residues

Table S6 reports per-residue mean RMSF for the 20 highest-plasticity binding-site residues in the SV2A common-basis TM backbone+C$\beta$ ensemble, ranked by max RMSF across the three trajectories. Binding-site residues are the SV2C contact residues within 5 Å of the bound ligand in the PSL-SV2C structure (Martin et al., manuscript in preparation), mapped to their SV2A equivalents via per-residue sequence alignment anchored on the conserved tryptophan cage (SV2A Trp300/Trp454/Trp666 $\leftrightarrow$ SV2C Trp286/Trp440/Trp651; $\Delta = 14$ through TM7, $\Delta = 15$ from TM8 onward, consistent with a single-residue indel in the TM7–TM8 loop). TM-helix assignments are from UniProt SV2A (Q7L0J3). The atom selection for per-residue RMSF was extended relative to the PCA figure (Figure 1) to include TM8 (SV2A residues 599–619; 1,954 atoms total vs. 1,804 for the PCA).

Table S6: Per-residue mean RMSF (Å) for the 20 highest-plasticity binding-site residues in the SV2A trajectories, ranked by max RMSF across the three conditions.

| SV2A | SV2C | TM | PRM (pH 5) | PRM (pH 7) | PSL |
|---|---|---|---|---|---|
| 608 | 593 | TM8 | 0.74 | 1.35 | 0.73 |
| 462 | 448 | TM7 | 0.82 | 1.22 | 0.56 |
| 461 | 447 | TM7 | 0.79 | 1.17 | 0.56 |
| 662 | 647 | TM10 | 0.65 | 1.03 | 0.67 |
| 297 | 283 | TM5 | 0.51 | 0.71 | 1.02 |
| 659 | 644 | TM10 | 0.79 | 1.01 | 0.78 |
| 183 | 169 | TM1 | 0.80 | 0.98 | 0.58 |
| 667 | 652 | TM10 | 0.54 | 0.77 | 0.95 |
| 694 | 679 | TM11 | 0.63 | 0.90 | 0.49 |
| 454 | 440 | TM7 | 0.63 | 0.88 | 0.63 |
| 689 | 674 | TM11 | 0.58 | 0.87 | 0.54 |
| 663 | 648 | TM10 | 0.70 | 0.86 | 0.69 |
| 686 | 671 | TM11 | 0.59 | 0.86 | 0.65 |
| 301 | 287 | TM5 | 0.62 | 0.83 | 0.71 |
| 693 | 678 | TM11 | 0.62 | 0.81 | 0.46 |
| 670 | 655 | TM10 | 0.50 | 0.74 | 0.76 |
| 690 | 675 | TM11 | 0.53 | 0.75 | 0.49 |
| 277 | 263 | TM4 | 0.60 | 0.71 | 0.59 |
| 666 | 651 | TM10 | 0.60 | 0.69 | 0.67 |
| 273 | 259 | TM4 | 0.48 | 0.65 | 0.58 |
